# Integration of multiple imaging platforms to uncover cardiac defects in adult zebrafish

**DOI:** 10.1101/2020.05.21.108548

**Authors:** Anabela Bensimon-Brito, Giulia L. M. Boezio, João Cardeira-da-Silva, Astrid Wietelmann, Christian S. M. Helker, Radhan Ramadass, Janett Piesker, Arno Nauerth, Clemens Mueller, Didier Y. R. Stainier

## Abstract

Mammalian models have been instrumental to investigate adult heart function and human disease. However, electrophysiological differences with human hearts and high costs emphasize the need for additional models. The zebrafish is a well-established genetic model to study cardiac development and function; however, analysis of cardiac phenotypes in adult specimens is particularly challenging as they are opaque. Here, we optimized and combined multiple imaging techniques including echocardiography, magnetic resonance imaging and micro-computed tomography to identify and analyze cardiac phenotypes in adult zebrafish. Using *alk5a/tgfbr1a* mutants as a case study, we observed morphological and functional cardiac defects, which were undetected with conventional approaches. Correlation analysis of multiple parameters revealed an association between hemodynamic defects and structural alterations of the heart, as observed clinically. Thus, we report a comprehensive and sensitive platform to identify otherwise indiscernible cardiac phenotypes in adult zebrafish, a model with clear advantages to study cardiac function and disease.

## Introduction

Cardiac diseases are a major cause of death worldwide (Rapezzi et al., 2013). The understanding of their pathophysiology has relied on the identification and analysis of genes triggering the initiation and progression of the cardiac defects. Several mammalian models including mice, rats, rabbits, pigs, sheep and dogs, have been used to investigate various cardiac diseases, each with its own benefits and shortcomings. For instance, rodent and human hearts differ in their electrophysiology and oxygen consumption (Nerbonne, 2004; Milani-Nejad et al., 2014), while large animals have considerable limitations in terms of cost, difficulty of handling, and genetic manipulation (Zaragoza et al., 2011; Milani-Nejad et al., 2014). Therefore, there is a need for additional animal models to investigate the mechanisms underlying cardiac diseases.

The zebrafish is a popular genetic model to study cardiac development, disease, and regeneration (Stainier et al., 1996; Stainier, 2001; Bakkers, 2011; Staudt et al., 2012). External fertilization and optical transparency during embryogenesis allow for *in vivo* analysis of cardiovascular development at single-cell resolution (Beis et al., 2006; Mickoleit et al., 2014). Previous studies have also highlighted the similarities between the mature human and zebrafish hearts, including comparable beating rates and action potential profiles (Sedmera et al., 2003; Kopp et al., 2005; Milan et al., 2006; Milan et al., 2008; Nemtsas et al., 2010; Vornanen et al., 2016; Ravens, 2018; Yan et al., 2020). Importantly, mutating zebrafish orthologs of human disease genes can accurately model cardiac disorders, as observed with mutations leading to arrhythmogenic, dilated and Titin-associated cardiomyopathies (Asimaki et al., 2014; Shih et al., 2015; Shih et al., 2016; Collins et al., 2019; Ding et al., 2019). Furthermore, unlike their mammalian counterparts, adult zebrafish hearts can regenerate, thereby providing a unique platform to understand mechanisms of cardiac regeneration (Poss et al., 2002; Gonzalez-Rosa et al., 2012; Gonzalez-Rosa et al., 2017). Together, these studies indicate that adult zebrafish have the unique potential to provide high-throughput, translatable scientific knowledge for pre-clinical and clinical research (Ding et al., 2013; Shih et al., 2015; Gut et al., 2017). However, morphological and functional characterization of adult zebrafish is particularly challenging as they are no longer optically transparent and too small for most conventional pre-clinical imaging platforms (Hein et al., 2015; Hur et al., 2017; Koth et al., 2017; Merrifield et al., 2017; Dvornikov et al., 2018). In addition, many cardiac disorders are associated with inherited genetic mutations (Kathiresan et al., 2012), often leading to variable phenotypic expressivity (Cooper et al., 2013). Therefore, to understand the cardiac phenotypes despite this variability between individuals, it is essential to extract all of the relevant information from a relatively large sample size, which is not easily done with mammalian models.

Due to the aforementioned imaging limitations, most studies of adult cardiac phenotypes in zebrafish rely on histological analyses of tissue sections which prevent three-dimensional (3D) representation of the morphological features or studies of cardiac function. In a clinical setting, diagnosing the onset and progression of cardiovascular diseases often requires a combination of multiple imaging analyses including echocardiography, magnetic resonance imaging (MRI) and computed tomography (CT) (Marwick et al., 2013; Fuchs et al., 2016; Petersen et al., 2017; Plana et al., 2018). Recently, researchers have adapted light-sheet microscopy (Packard et al., 2017), MRI (Koth et al., 2017; Merrifield et al., 2017) and micro-computed tomography (μ-CT) (Descamps et al., 2014; Weinhardt et al., 2018; Ding et al., 2019) to define the morphology of zebrafish organs. Others have used echocardiography to characterize the functional impact of cardiac disease and regeneration in adult zebrafish (Ho et al., 2002; Sun et al., 2008; Parente et al., 2013; González-Rosa et al., 2014; Hein et al., 2015; Huang et al., 2015; Wilson et al., 2015; Ernens et al., 2016; Lee et al., 2016; Wang et al., 2017; Benslimane et al., 2019). However, a systematic workflow to study cardiac phenomics in adult zebrafish has yet to emerge.

Here, we established, optimized, and combined functional analyses of the adult zebrafish heart using echocardiography and MRI with a morphological characterization of the cardiac compartments and adjacent vessels using μ-CT. As a proof-of-concept, we examined *alk5a/tgfbr1* mutants as *TGFBR1* genetic variants in humans have been associated with aortic aneurysms and diseases of the great vessels, the diagnosis of which is most often recognized in young adults (Loeys et al., 2005; Loeys et al., 2006; Matyas et al., 2006; Weerakkody et al., 2018; Camerota et al., 2019). We show that the use of this multimodal preclinical imaging strategy allows for a robust dissection of adult zebrafish cardiac phenotypes, even in the presence of high variability between samples of the same genotype.

## Results

### Selecting a mutant to test the sensitivity of our imaging platform

In order to test the power of multiple imaging techniques to detect subtle adult phenotypes, we first needed to select a suitable model. Many homozygous mutant zebrafish exhibiting cardiac defects at early developmental stages have been reported (Sehnert et al., 2002; Miura et al., 2011; Tu et al., 2012; Wilkinson et al., 2014; Grant et al., 2017), and some animals heterozygous for mutations display subtle defects in adult stages (Ahlberg et al., 2018; Huttner et al., 2018). Other zebrafish mutants do not display embryonic phenotypes but exhibit pericardial edema at adult stages (Guerra et al., 2018), indicating cardiac function defects. We elected to investigate *alk5a* mutants (Figure 1A), a third type of mutant, i.e., one that does not exhibit embryonic phenotypes or pericardial edema at adult stages but can potentially develop cardiac phenotypes. *alk5a*; *alk5b* double mutants exhibit severe pericardial edema at early developmental stages, due to a defective outflow tract (OFT) (Boezio et al., 2020). In contrast, *alk5a* single mutant larvae do not exhibit any obvious defects (Figure 1 - figure supplement 1 A, B), including in the cardiovascular system (Figure 1 - figure supplement 1 C-D’), and survive to adulthood. Similarly, *alk5a*^*-/-*^ animals at 9 months post fertilization (mpf) display no gross morphological defects or pericardial edema (Figure 1B, C). However, analysis of dissected hearts including histological sections suggested that some *alk5a* ^*-/-*^ adult fish have an expanded OFT lumen when compared to wild type (WT; Figure 1D-I). The levels of Elastin2, a major component of the ECM in the cardiac OFT, appear to be reduced in the *alk5a*^*-/-*^ OFT wall (Figure 1F, G). However, as histological sections do not yield a 3D representation of morphological features, it was not possible to conclude whether the OFT lumen was indeed dilated in *alk5a* mutants. Moreover, due to the opacity of adult zebrafish, one cannot use live microscopy to investigate the functional consequences of morphological cardiac defects. Therefore, we aimed to adapt and optimize cardiovascular preclinical imaging techniques to collect multiple cardiac parameters (Table 1) and analyze adult zebrafish phenotypes. We randomly selected 10 WT and 12 *alk5a* mutant zebrafish, ranging from 6 to 15 mpf (Figure 1J, Table 2), and tracked each individual throughout all the analyses. To account for any major morphological differences between the two groups, we measured standard length (Figure 1K) and body volume (Figure 1L) and observed no significant differences between WT and mutant groups.

**Table 1.** Description and abbreviation terms for each parameter.

| Parameter | Abbreviation | Description |
| --- | --- | --- |
| Standard length | Std. length | Length from the anterior tip of the snout to the anterior part of the caudal fin |
| Body volume | Body vol. | Three-dimensional space occupied by the fish body |
| Outflow tract velocity time integral | OFT VTI | Area within the Doppler curve (integral). It indicates how far blood travels during the flow period and it is a measure of cardiac systolic function and cardiac output. |
| Outflow tract mean velocity | OFT mean velocity | Average of the instantaneous velocity from the outflow curve for each heartbeat, measured from all heart cycles across 10 seconds |
| Outflow tract mean gradient | OFT mean gradient | Average of the instantaneous pressure gradients over the entire outflow Doppler curve, measured from all heart cycles across 10 seconds |
| Outflow tract velocity peak velocity | OFT peak velocity | Maximum velocity of the blood passing from the ventricle to the outflow tract; $V_{\max}$ |
| Outflow tract velocity peak gradient | OFT peak gradient | Pressure gradient calculated from the peak velocity with the Bernoulli equation: $4(V_{\max})^2$ |
| Ejection time | Ejection time | Period of blood flow from ventricle to the outflow tract |
| Heart rate | Heart rate | The speed at which the heart beats measured in beats per minute |
| Area aortic flow | Area aortic flow | Amount of blood (in a 2D plane) detected in the aorta |
| Area inflow | Area inflow | Amount of blood (in a 2D plane) going from the atrium to the ventricle |
| Area outflow | Area outflow | Amount of blood (in a 2D plane) going from the ventricle to the outflow tract |
| Regurgitation fraction in the atrioventricular canal | Regurg. fraction AVC | Relative amount of blood (in a 2D plane) that returns to the atrium during ventricular contraction (systole); inflow area/retrograde flow area |
| Regurgitation fraction in the outflow tract | Regurg. fraction OFT | Relative amount of blood (in a 2D plane) that returns to the ventricle after ventricular contraction (systole); outflow area/retrograde flow area |
| Percentage ventricular luminal expansion | % Vent. lum. exp. | Lumen displacement between minimum and maximum ventricular contraction in a heart cycle |
| Percentage ventricular outer wall expansion | % Vent. outer wall exp. | Outer wall displacement between minimum and maximum ventricular contraction in a heart cycle |
| Percentage outflow tract luminal expansion | % OFT lum. exp. | Lumen displacement between minimum and maximum outflow tract expansion in a heart cycle |
| Percentage outflow tract outer wall expansion | % OFT outer wall exp. | Outer wall displacement between minimum and maximum outflow tract expansion in a heart cycle |
| Heart volume | Heart vol. | Three-dimensional space occupied by the heart – calculated by the sum of the volumes of each cardiac compartment |
| Ventricular volume | Vent. vol. | Three-dimensional space occupied by the ventricle – including the lumen |
| Percentage ventricular volume | % Vent. vol. | Relative volume of the ventricle in relation to the entire heart |
| Atrial volume | Atrial vol. | Three-dimensional space occupied by the atrium – including the lumen |
| Percentage atrial volume | % Atrial vol. | Relative volume of the atrium in relation to the entire heart |
| Outflow tract volume | OFT vol. | Three-dimensional space occupied by the outflow tract – including the lumen |
| Percentage of outflow tract volume | % OFT vol. | Relative volume of the outflow tract in relation to the entire heart |
| Outflow tract luminal volume | OFT lum. vol. | Three-dimensional space occupied by the outflow tract lumen |
| Percentage outflow tract luminal volume | % OFT lum. vol. | Relative volume of the outflow tract lumen in relation to the entire outflow tract |
| Outflow tract wall volume | OFT wall vol. | Three-dimensional space occupied by the outflow tract wall; OFT volume - OFT lumen volume |
| Aortic diameter | Aortic diam. | Maximum diameter of the aorta measured in the most proximal linear portion of the aorta |
| Aortic volume | Aortic vol. | Three-dimensional space occupied by the aorta anteriorly to the heart |

**Table 2.**
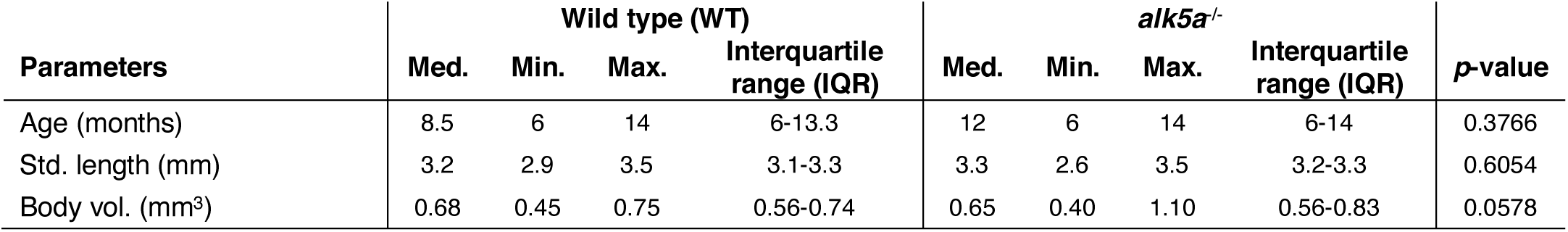
Biological features.

| Parameters | Wild type (WT) |  |  |  | *alk5a*<sup>-/-</sup> |  |  |  | *p*-value |
| --- | --- | --- | --- | --- | --- | --- | --- | --- | --- |
|  | Med. | Min. | Max. | Interquartile range (IQR) | Med. | Min. | Max. | Interquartile range (IQR) |  |
| Age (months) | 8.5 | 6 | 14 | 6-13.3 | 12 | 6 | 14 | 6-14 | 0.3766 |
| Std. length (mm) | 3.2 | 2.9 | 3.5 | 3.1-3.3 | 3.3 | 2.6 | 3.5 | 3.2-3.3 | 0.6054 |
| Body vol. (mm<sup>3</sup>) | 0.68 | 0.45 | 0.75 | 0.56-0.74 | 0.65 | 0.40 | 1.10 | 0.56-0.83 | 0.0578 |

**Figure 1.**
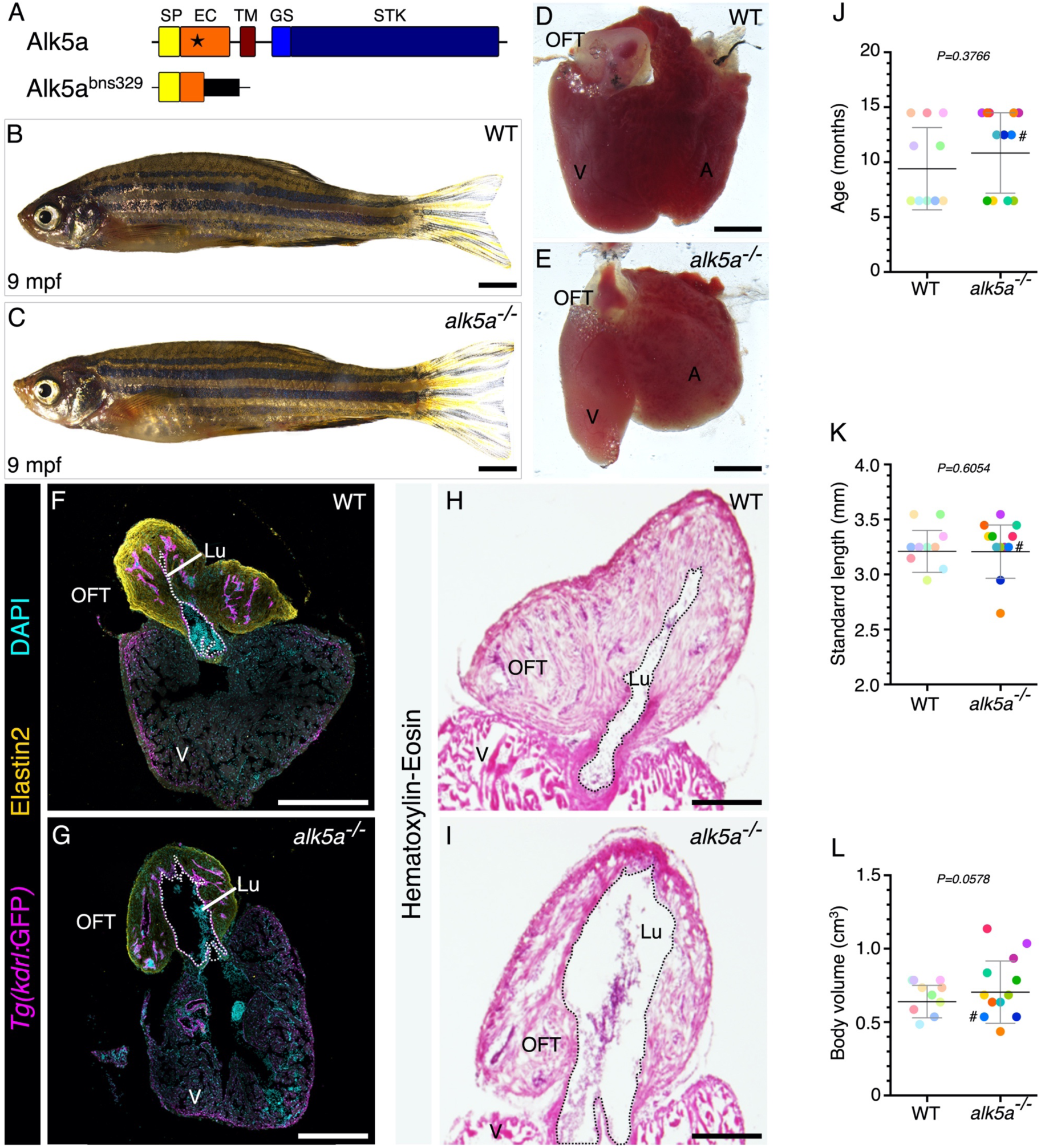
*alk5a* mutant adult zebrafish display variable cardiac phenotypes without gross morphological defects. (A) Schematic of Alk5a WT and mutant proteins depicting each domain and the site of the mutation (star). (B, C) Brightfield images of 9 mpf WT (B) and *alk5a*^*-/-*^ (C) zebrafish. (D, E) Brightfield images of *alk5a*^*-/-*^ hearts (E) which occasionally exhibit a dilatedoutflow tract (OFT) lumen compared to wild type (D). (F-I) Cryosections of WT (F, H) and *alk5a*^*-/-*^ (G, I) hearts immunostained for *Tg(kdrl:*eGFP) expression (endothelial cells) and Elastin2 (F, G), and stained for hematoxylin-eosin (H, I) showing the expanded OFT lumen (Lu, dashed line) in *alk5a* ^*-/-*^ mutants. (J-L) Quantification of age (G), standard length (H), and body volume (I) of WT and *alk5a*^*-/-*^ zebrafish used in the subsequent analyses. Plots show the values for each individual and the mean ± SD; *P*-values were determined by unpaired *t*-test (J, L) or Mann-Whitney test (K). The color of each dot refers to the same fish across all graphs. The number symbol refers to the individual fish mentioned in the text. SP - signal peptide, EC – extracellular, TM -transmembrane, GS -glycine-serine rich, STK - serine-threonine kinase, A- atrium, V- ventricle. Scale bars: 2 mm (B, C), 200 μm (D, E, H, I), 400 μm (F, G).

### Echocardiography reveals variable hemodynamic defects in *alk5a* mutants

Echocardiography, the fastest (approximately 5 minutes per fish) and least invasive imaging technique, records hemodynamic parameters in live animals (Scherrer-Crosbie et al., 2008; Moran et al., 2013; Gaspar et al., 2018; Wang et al., 2018). We analyzed cardiac function from sagittal views of the heart (Figure 2 - figure supplement 1 A), and through Doppler echocardiography were able to assess a range of features including the direction, amount, and velocity of blood flow (Figures 2 A-D and Figure 2 - figure supplement 1 B-C’, Table 3). Due to the intrinsic variability between individuals, the heart rate ranged from 58 to 143 beats per minute (bpm), with no statistically significant difference between the two genotypes (Figure 2A). We also determined the area of blood inflow and outflow on a two-dimensional (2D) plane and noticed that while most fish exhibited unidirectional inflow (red; 7/10 in WT and 6/12 in *alk5a*^*-/-*^; Figure 2C, E, Video 1) and outflow (blue; 8/10 in WT and 5/12 in *alk5a*^*-/-*^; Figure 2D, E’, Video 1), some mutant fish (7/12) also presented blood flow regurgitation in the atrioventricular canal (AVC; Figure 2C, F’, G, Videos 2 and 3) and/or in the OFT canal (Figure 2 D, G’, Video 3). In addition, while most fish exhibited a flow signal exclusively in the heart, a few displayed a signal rostral to the OFT, in the region of the ventral aorta, hereafter referred to as aortic flow (Video 2). The presence of aortic flow was detected more frequently in *alk5a*^*-/-*^ fish (1/10 in WT and 4/12 in *alk5a*^*-/-*^; Figure 2B, F), suggesting an increase in blood flow in their aorta. Nonetheless, the relative inflow and outflow areas in a 2D plane, as well as most hemodynamic parameters including blood flow velocity and ejection time were highly variable and not significantly different between WT and mutant fish (Figure 2 - figure supplement 1 D-K). In contrast, by determining several parameters in each fish, we identified one mutant (indicated by # in all plots) with a large inflow area (Figure 2 - figure supplement 1 J) that also displayed severe blood regurgitation in the AVC and OFT (Figure 2 C, D), as well as reduced blood flow velocities through the OFT (Figure 2 - figure supplement 1 D, E, G).

**Table 3.** Echocardiography parameters.

| Parameters | Wild type (WT) |  |  |  | *alk5a*<sup>-/-</sup> |  |  |  | *p*-value |
| --- | --- | --- | --- | --- | --- | --- | --- | --- | --- |
|  | Med. | Min. | Max. | Interquartile range (IQR) | Med. | Min. | Max. | Interquartile range (IQR) |  |
| OFT VTI (mm) | 12.3 | 6.9 | 28.5 | 11.2-19.4 | 11.8 | 1.8 | 20.3 | 9.2-16.4 | 0.2997 |
| OFT mean velocity (mm/s) | 77.4 | 44.7 | 132.0 | 64.4-94.9 | 75.2 | 17.6 | 123.5 | 59.6-95.3 | 0.5757 |
| OFT mean gradient (mm Hg) | 0.024 | 0.008 | 0.070 | 0.017-0.036 | 0.022 | 0.001 | 0.062 | 0.014-0.037 | 0.5903 |
| OFT peak velocity (mm/s) | 110.4 | 59.7 | 186.1 | 86.6-128.9 | 98.9 | 25.6 | 171.8 | 82.4-129.4 | 0.5278 |
| OFT peak gradient (mm Hg) | 0.050 | 0.014 | 0.139 | 0.030-0.067 | 0.040 | 0.003 | 0.119 | 0.027-0.066 | 0.5229 |
| Ejection Time (s) | 0.182 | 0.136 | 0.246 | 0.159-0.200 | 0.157 | 0.103 | 0.237 | 0.137-0.181 | 0.1947 |
| Heart rate (bpm) | 107.8 | 58.4 | 131.5 | 87.4-118.9 | 98.4 | 61.0 | 143.5 | 84.8-112.3 | 0.8138 |
| Area aortic flow (x10<sup>3</sup> mm<sup>2</sup>) | 0 | 0 | 0.67 | 0 | 0 | 0 | 3.79 | 0-0.161 | 0.1729 |
| Area inflow (x10<sup>4</sup> AU) | 0.81 | 0.45 | 1.24 | 0.67-0.97 | 0.52 | 0.04 | 3.69 | 0.38-0.82 | 0.1583 |
| Area outflow (x10<sup>3</sup> AU) | 6.40 | 3.27 | 11.75 | 5.22-6.89 | 4.12 | 0.56 | 10.09 | 3.47-5.90 | 0.1447 |
| Regurg. fraction AVC (%) | 0 | 0 | 11.2 | 0-1.2 | 2.6 | 0 | 84.0 | 0-27.0 | 0.1732 |
| Regurg. fraction OFT (%) | 0 | 0 | 24.4 | 0 | 6.0 | 0 | 55.1 | 0-16.0 | 0.0847 |

**Figure 2.**
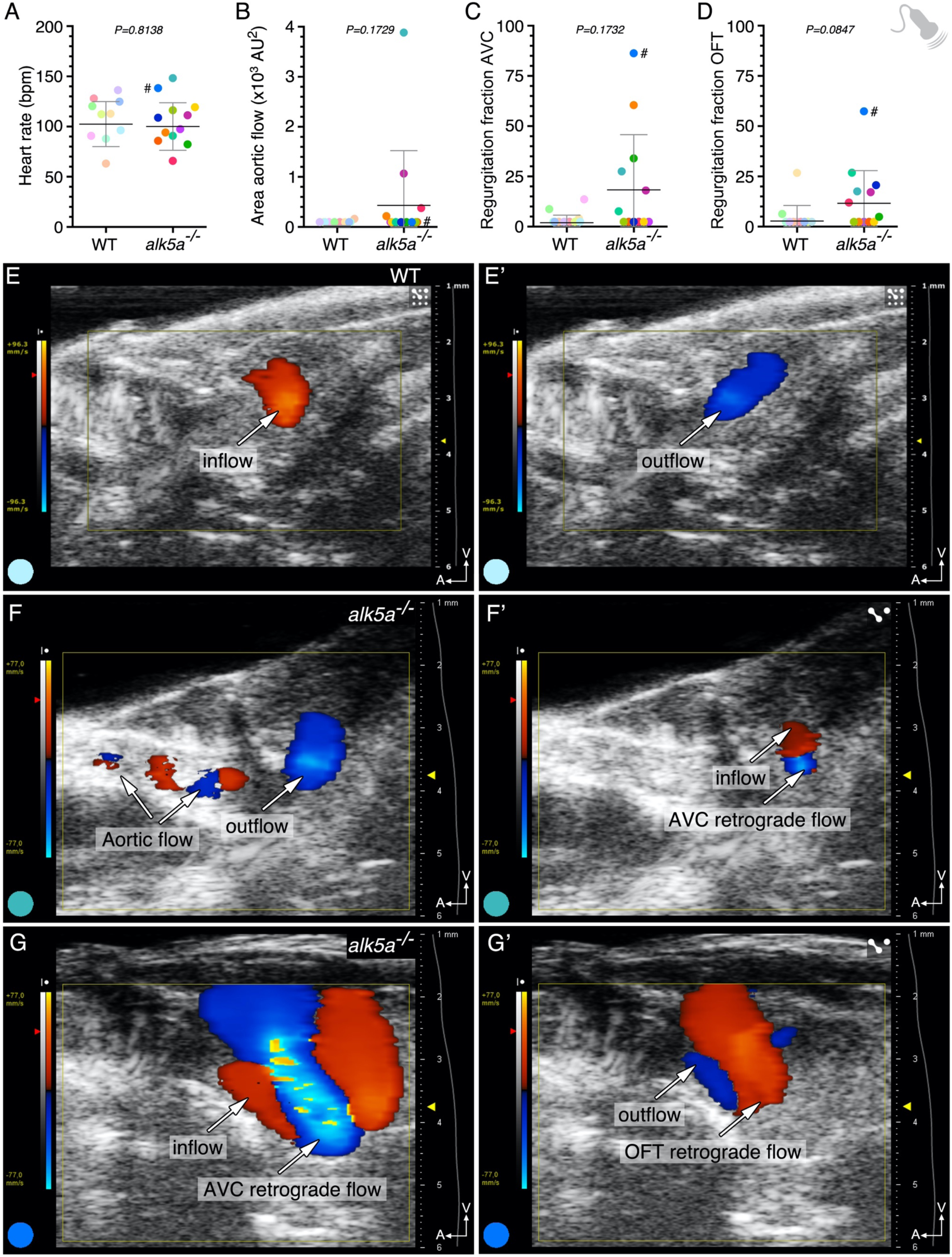
*alk5a* mutant adults display variable hemodynamic defects. (A-D) Parameter quantification obtained with echocardiography analyses of WT and *alk5a*^*-/-*^ adult zebrafish, including heart rate (A), area of the aortic flow (B) and regurgitation fraction in the AV (C) and OFT canals (D). Plots show the values for each individual and the mean ± SD; *P*-values were determined by unpaired *t-*test (A) or Mann Whitney test (B-D). (E, E’) WT zebrafish exhibit unidirectional blood inflow (red, E) and outflow (blue, E’) without signs of regurgitation. (F-G’) Examples of *alk5a*^*-/-*^ zebrafish exhibiting a detectable aortic flow (F) and retrograde blood flow (F’-G’). The color of each dot refers to the same fish across all graphs and images. The number symbol refers to the individual fish mentioned in the text.

In summary, despite the great variability obtained with echocardiography, this technique provides important information on cardiac function, particularly regarding hemodynamics in the heart and adjacent vessels.

### MRI analyses of beating hearts reveal expansion of most *alk5a* mutant outflow tracts

To achieve higher resolution of cardiac morphology and performance, we used MRI to study the same 22 zebrafish. MRI is the gold standard technique for clinical analyses of cardiac morphology and function due to its precision and high reproducibility (Ambale-Venkatesh et al., 2015; Kawel-Boehm et al., 2015; Axel et al., 2016; Captur et al., 2016; Dastidar et al., 2016; de Roos, 2019). Although MRI has been increasingly used in small animal models including mice (Nahrendorf et al., 2003; Price et al., 2010; Braig et al., 2017), in zebrafish it has been restricted to *ex-vivo* samples and static images of the regenerating heart (Kabli et al., 2010; Ullmann et al., 2010; Koth et al., 2017; Merrifield et al., 2017). In rodent models, MRI permits the virtual reconstruction of beating hearts (Echteld et al., 2006; Bovens et al., 2011), but this type of analysis is challenging in adult zebrafish due to the small size of their heart. Hence, we optimized cardiac MRI for adult zebrafish to precisely assess cardiac function in live animals (see Materials and Methods).

We imaged the heart in two different planes to obtain coronal views of the OFT and ventricle and to observe the expansion of these compartments during the cardiac cycle (Figure 3 - figure supplement 1 A-B’). Additionally, using self-gating signals (Echteld et al., 2006; Bovens et al., 2011), we were able to monitor the cardiac rate of the animal (Figure 3 - figure supplement 1 C-C’’) to accurately reconstruct the cardiac cycle. Then, we calculated the percentage of the cardiac chambers’ expansion, measuring both the outer wall and the lumen (Figures 3A-D’’ and Figure 3 - figure supplement 1 D-G’’, Table 4). We found that the expansion of the ventricular outer wall and of the ventricular lumen during the cardiac cycle were comparable between WT and *alk5a*^*-/-*^ fish (Figure 3 - figure supplement 1 D, E, Videos 4 and 5). In contrast, we observed that the mutants displayed a severe and consistent increase in OFT luminal dilation (Figure 3B, C-D’’, Videos 6 and 7), whereas the OFT outer wall expansion was comparable to that of WT (Figure 3A, C-D’’). Also, when analyzing the results for the mutant with severe hemodynamic defects detected with echocardiography (#), we observed that it had one of the highest OFT luminal expansion levels (Figures 3B).

**Table 4.** MRI parameters.

| Parameters | Wild type (WT) |  |  |  | *alk5a*<sup>-/-</sup> |  |  |  | *p*-value |
| --- | --- | --- | --- | --- | --- | --- | --- | --- | --- |
|  | Med. | Min. | Max. | Interquartile range (IQR) | Med. | Min. | Max. | Interquartile range (IQR) |  |
| Vent. lum. exp. (%) | 27.3 | 8.0 | 59.0 | 17.6-31.8 | 28.3 | 17.5 | 63.0 | 20.9-36.4 | 0.5277 |
| Vent. outer wall exp. (%) | 9.0 | 2.5 | 22.0 | 3.1-15.0 | 6.5 | 1.0 | 20.5 | 4.4-12.6 | 0.8107 |
| OFT lum. exp. (%) | 10.8 | 7.0 | 18.0 | 9.8-13.0 | 36.0 | 12.0 | 41.0 | 33.0-38.3 | **<0.0001** |
| OFT outer wall exp. (%) | 6.3 | 4.0 | 10.0 | 4.4-7.5 | 6.8 | 3.0 | 12.0 | 5.1-8.1 | 0.6395 |

**Figure 3.**
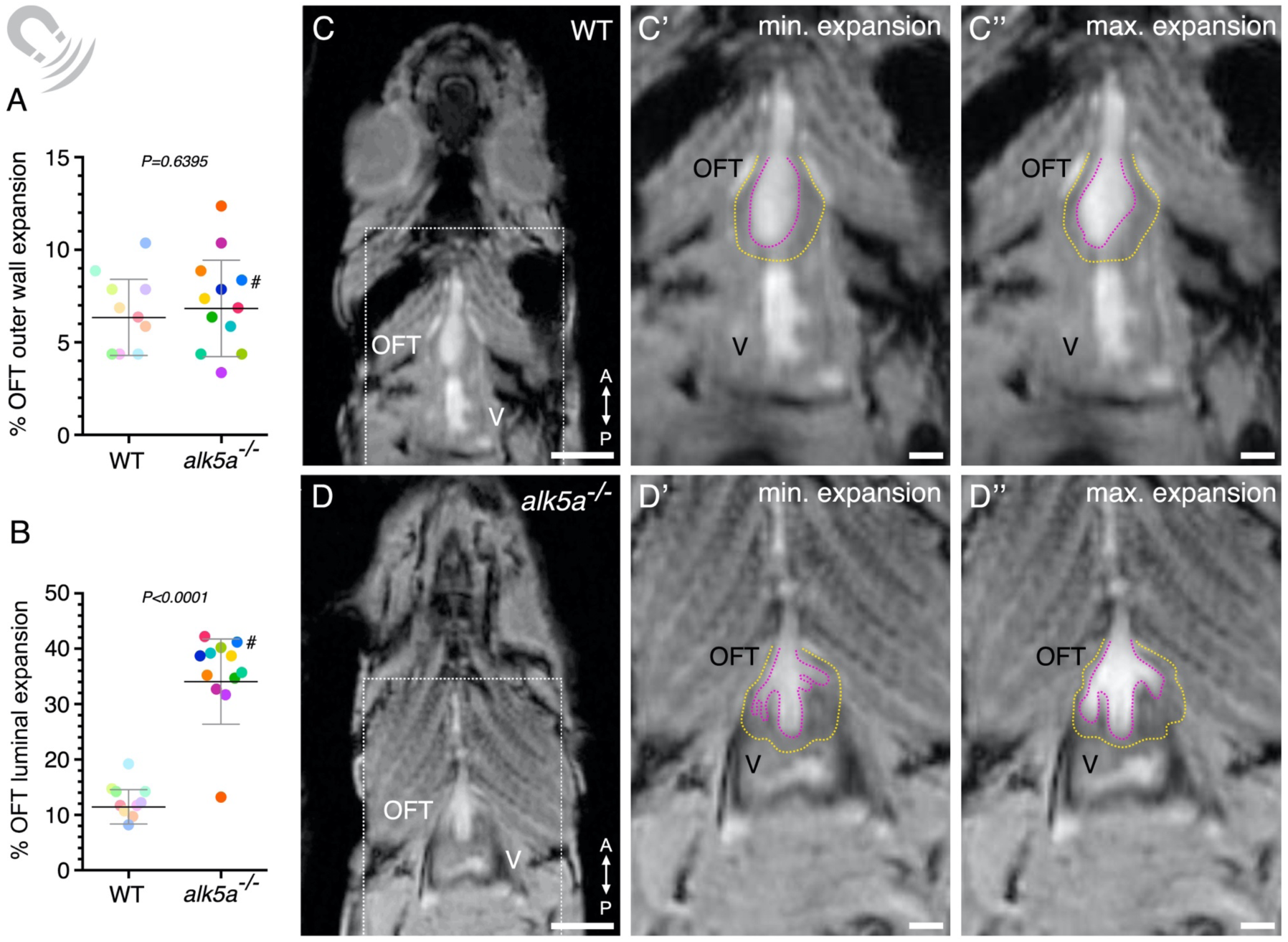
MRI analyses of beating hearts reveal increased OFT luminal dilation in *alk5a* mutants. (A, B) Percentage of OFT outer wall (A) and luminal (B) expansion in WT and *alk5a*^*-/-*^ adult zebrafish. Plots show the values for each individual and the mean ± SD; *P*-values were determined by unpaired *t*-test (A) or Mann-Whitney test (B). (C-D’’) Single frames of MRI cines of WT (C-C’’) and *alk5a*^*-/-*^ (D-D’’) zebrafish in coronal view. Boxed area is shown in C’, D’ (minimum expansion OFT) and C’’, D’’ (maximum expansion OFT). Magenta dashed line, OFT lumen; yellow dashed line, OFT outer wall. The color of each dot refers to the same fish across all graphs and images. The number symbol refers to the individual fish mentioned in the text. V- ventricle, OFT- outflow tract. Scale bars: 1.5 mm (C, D), 500 μm (C’, C’’, D’, D’’).

In summary, by optimizing MRI imaging of beating adult zebrafish hearts, we were able to obtain unique and valuable data on cardiac performance *in vivo*. Moreover, we identified the OFT as the main cardiac compartment affected in *alk5a* mutants.

### micro-CT analyses reveal outflow tract defects in *alk5a* mutants

X-ray based *ex vivo* CT is considered the most powerful technique to provide high spatial resolution of small structures (Liguori et al., 2015; Larsson et al., 2016; Mortensen et al., 2018; Litmanovich et al., 2019). Here we use μ**-**CT, which is capable of volumetric analysis with a voxel size <20 μm (Holdsworth et al., 2002; Clark et al., 2014; Senter-Zapata et al., 2016) to characterize the 3D morphology of the cardiac compartments and adjacent vessels. Due to the exposure to radiation, the requirement for a contrast agent, and for long imaging periods to acquire high-resolution images, we performed this imaging on fixed samples. Individual X-ray projections were used to reconstruct cross-section images, providing views of the tissues in multiple planes (Figure 4 - figure supplement 1 A-A’’).

After manual segmentation and volumetric surface rendering (Figure 4A-D’’’’, Videos 8-11), we could determine the volume of each cardiac compartment, as well as the aortic diameter and volume (Figures 4E-J and Figure 4 - figure supplement 1 B-G, Table 5, Videos 12 and 13). Consistent with the measurements obtained with the other techniques, we did not observe any statistically significant differences between WT and *alk5a*^*-/-*^ fish regarding the volume of the entire heart (Figure 4E), atrium (Figures 4F, Figure 4 - figure supplement 1 D), or ventricle (Figure 4 - figure supplement 1 B, C). Moreover, while the total volume of the OFT appeared to be lower in *alk5a*^*-/-*^ fish, this difference was not statistically significant (Figures 4G and Figure 4 - figure supplement 1 E). However, the mutants displayed a statistically significant increase in the OFT luminal volume (relative to the OFT volume; Figure 4H), and a decrease in the OFT wall volume (Figure 4I). Despite the observation of aortic flow in mutant fish suggesting higher aortic volumes, *alk5a*^*-/-*^ zebrafish only exhibited a slight increase in aortic diameter compared to WT (Figure 4C’’’’, D’’’’, J, Videos 14 and 15), while displaying an unaltered volume (Figure 4 - figure supplement 1 G).

**Table 5.** micro-CT parameters.

| Parameters | Wild type (WT) |  |  |  | *alk5a*<sup>-/-</sup> |  |  |  | *p*-value |
| --- | --- | --- | --- | --- | --- | --- | --- | --- | --- |
|  | Med. | Min. | Max. | Interquartile range (IQR) | Med. | Min. | Max. | Interquartile range (IQR) |  |
| Heart vol. (x10<sup>9</sup> μm<sup>3</sup>) | 3.11 | 2.18 | 4.20 | 2.49-3.84 | 2.95 | 1.67 | 7.67 | 2.74-3.62 | 0.9022 |
| Vent. vol. (x10<sup>8</sup> μm<sup>3</sup>) | 1.40 | 1.00 | 2.00 | 1.19-1.75 | 1.41 | 6.07 | 1.99 | 9.28-10.53 | 0.3142 |
| Vent. vol. (%) | 46.0 | 34.2 | 61.0 | 43.3-49.5 | 46.0 | 8.1 | 53.5 | 41.7-49.6 | 0.9094 |
| Atrial vol. (x10<sup>9</sup> μm<sup>3</sup>) | 1.33 | 9.91 | 2.41 | 1.08-2.01 | 1.38 | 9.68 | 6.96 | 1.12-1.78 | 0.9370 |
| Atrial vol. (%) | 48.3 | 30.2 | 61.4 | 43.4-51.0 | 48.0 | 39.4 | 90.7 | 43.8-54.4 | 0.7354 |
| OFT vol. (x10<sup>8</sup> μm<sup>3</sup>) | 1.78 | 1.15 | 2.90 | 1.52-2.27 | 1.82 | 6.02 | 2.13 | 1.06-1.86 | 0.1074 |
| OFT vol. (%) | 6.2 | 4.4 | 8.8 | 4.9-7.1 | 5.4 | 1.3 | 7.2 | 3.9-6.6 | 0.1328 |
| OFT lum. vol. (x10<sup>7</sup> μm<sup>3</sup>) | 0.78 | 0.16 | 2.30 | 0.59-1.50 | 2.19 | 0.25 | 4.95 | 1.08-2.90 | 0.1328 |
| OFT lum. vol. (%) | 4.0 | 1.1 | 10.7 | 0.3-0.7 | 11.6 | 4.2 | 51.5 | 7.5-14.9 | **0.0034** |
| OFT wall vol. (x10<sup>8</sup> μm<sup>3</sup>) | 1.72 | 1.13 | 2.69 | 1.46-2.09 | 1.59 | 0.47 | 1.83 | 0.91-1.71 | **0.0284** |
| Aortic diam. (μm) | 193.0 | 138.0 | 253.0 | 157.8-199.0 | 196.0 | 148.0 | 300.0 | 169.5-258.3 | 0.1724 |
| Aortic vol. (x10<sup>7</sup> μm<sup>3</sup>) | 4.06 | 2.17 | 7.73 | 3.31-6.46 | 4.09 | 1.55 | 9.64 | 3.18-6.63 | 0.8638 |

**Figure 4.**
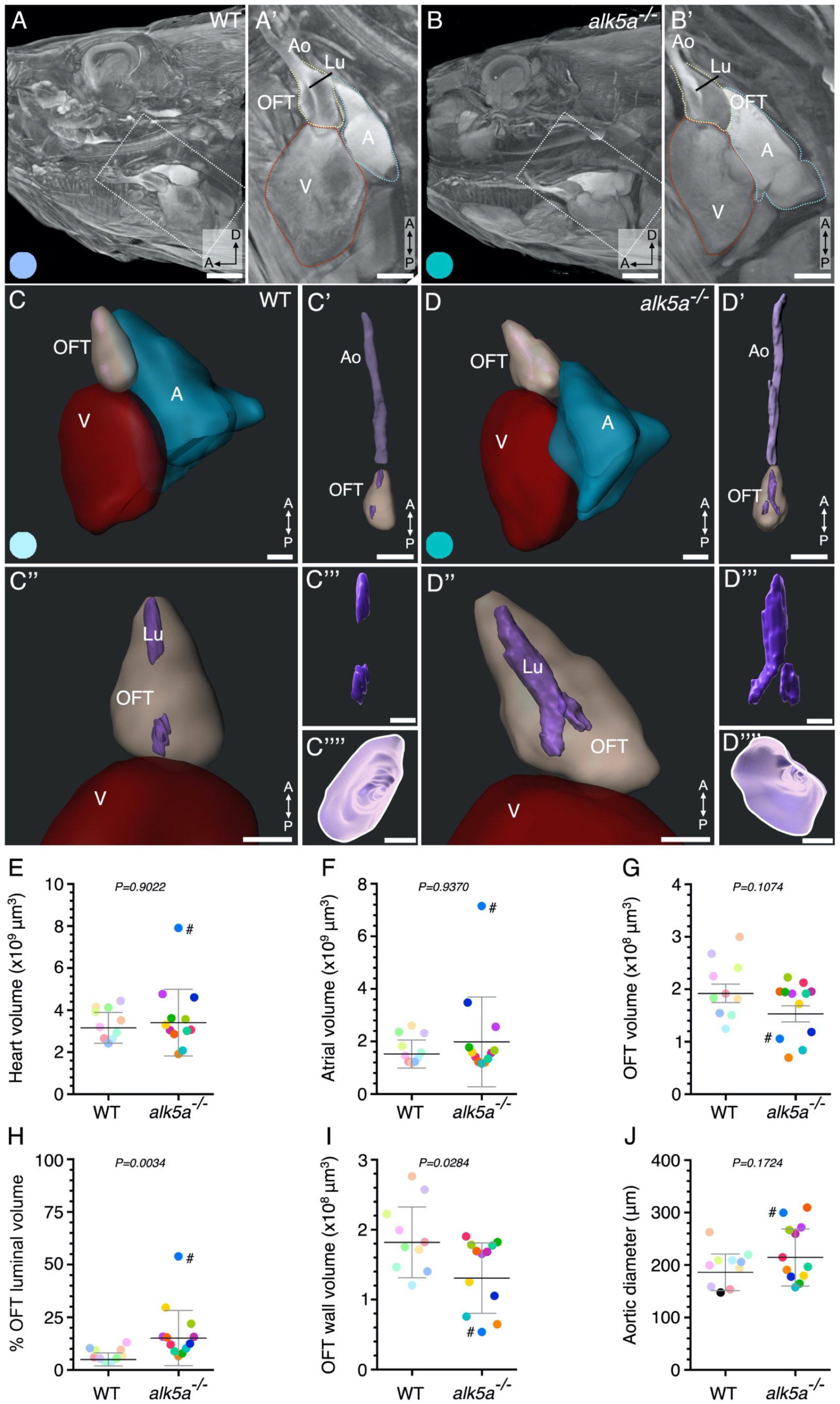
μ-CT analyses of cardiovascular morphology in *alk5a* mutants reveal OFT defects. (A-B) μ-CT scans showing a sagittal plane of the anterior region of WT and *alk5a*^*-/-*^ adult zebrafish. Boxed area is shown in A’ and B’. Dashed lines outline cardiac compartments. (C-D’’’’) 3D reconstructions of the cardiac compartments in WT (C-C’’’’) and *alk5a*^*-/-*^ (D-D’’’’) zebrafish, including the aorta (C’, D’), and OFT lumen (Lu; C’’’, D’’’). (C’’’’, D’’’’) 3D reconstructions of the aortic opening (white line) as seen from the OFT. (E-J) Quantification of morphological parameters for each cardiac compartment in WT and *alk5a*^*-/-*^ zebrafish. Plots show the values for each individual and the mean ± SD; *P*-values were determined by unpaired *t-*test (G-J) or Mann-Whitney test (E, F). The color of each dot refers to the same fish across all graphs and images. The number symbol refers to the individual fish mentioned in the text. A – atrium, Ao – aorta, V – ventricle, OFT-outflow tract. Scale bars: 1 mm (A, B), 500 μm (A’, B’), 300 μm (C-D’), 400 μm (C’’-D’’’’).

It is important to note that the μ-CT analyses provided sufficient resolution to visualize smaller structures such as the ventricular trabeculae (Figures 4A-B’ and Figure 4 - figure supplement 1 A’’) and the atrioventricular valve (Figure 4 B’). Moreover, by tracking individual fish with the different techniques, we observed that the animal displaying the most severe functional phenotype as assessed by echocardiography and MRI (#) also presented severe morphological defects when analyzed with μ**-**CT. In particular, this individual exhibited the largest OFT luminal volume (Figure 4H) and one of the largest aortic diameters (Figure 4J), but the smallest OFT wall volume (Figure 4I). This fish also displayed an abnormally large atrial volume (Figure 4F), in line with the high regurgitation fraction in the AVC identified by echocardiography.

Thus, with the help of μ**-**CT imaging, we were able to obtain precise structural information about all the cardiac compartments and further resolve the OFT defects identified using the other platforms.

### Correlation analysis between all parameters reveals previously undetected phenotypes in *alk5a* mutants

From the 32 parameters measured with the different imaging platforms, only 3 showed a *p*<0.05 between WT and mutant samples after bulk analysis. The lack of statistical significance in group comparisons is due to the intrinsic variability between individuals of the same genotype. However, when one follows a single individual (e.g., #), it becomes clear that a higher severity in functional phenotypes is accompanied by stronger morphological alterations. Therefore, we analyzed how variables were linked by determining the pairwise Pearson’s correlation coefficients across all the WT and *alk5a* mutant animals, herein represented as correlograms (WT, Figure 5A, and *alk5a*^*-/-*^, Figure 5B).

**Figure 5.**
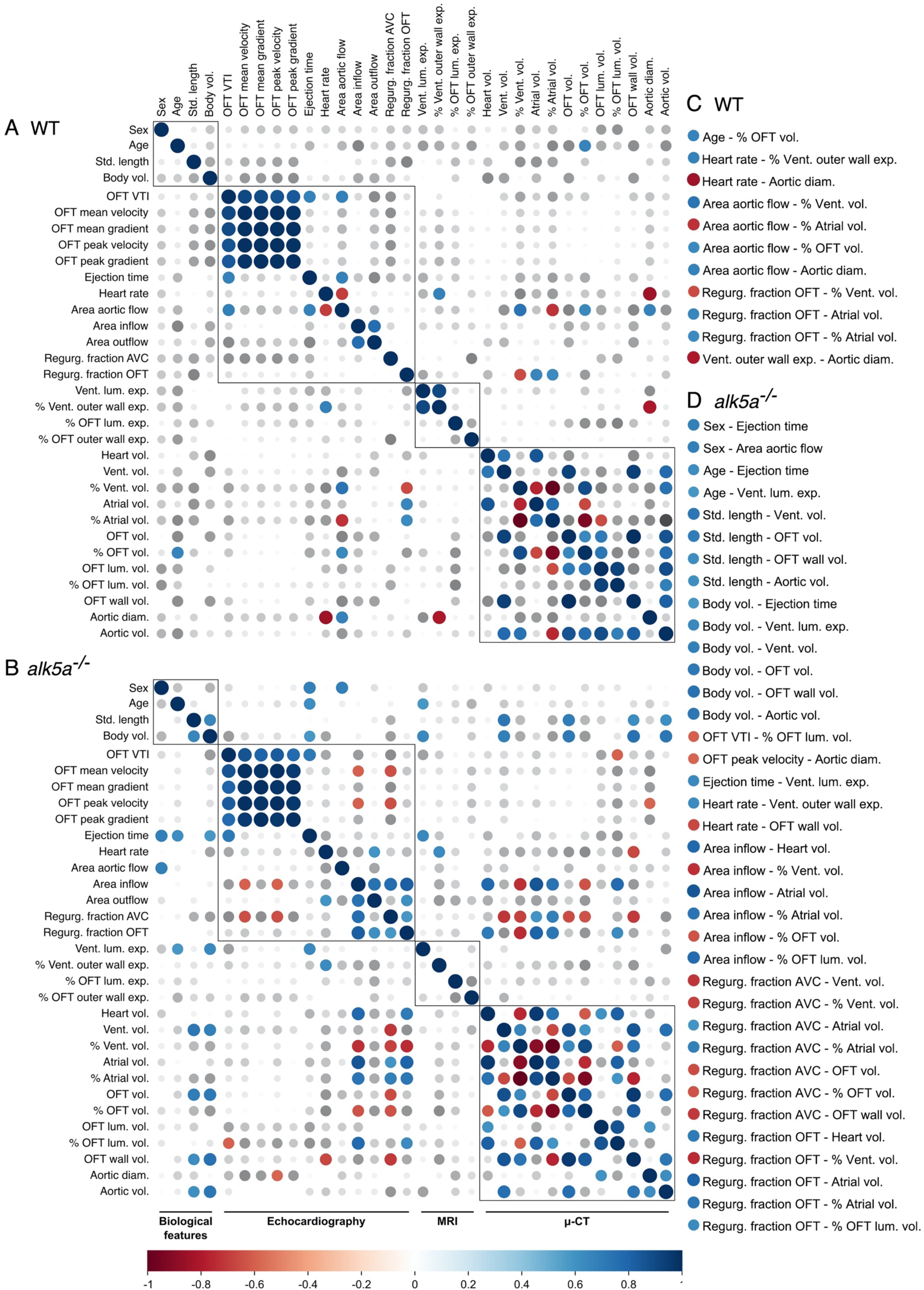
Correlation analysis of all the measured parameters reveals previously undetected cardiovascular phenotypes. (A, B) Correlograms of all the parameters analyzed including biological features, echocardiography, MRI and µ-CT analyses across all the WT (A) and *alk5a*^*-/-*^ (B) zebrafish. Positive and negative correlations are shown in blue and red, respectively; dot size represents Pearson’s correlation coefficient. (C, D) Significant correlations (*P-*value < 0.05) in WT (C) and *alk5a*^*-/-*^ (D) zebrafish.

As expected, we observed strong correlations between measurements obtained within the same platform, but also several interesting inter-platform correlations. As seen in humans whose cardiac parameters strongly correlate with gender, age and ethnicity (Lancellotti, 2014; Caballero et al., 2015; Kawel-Boehm et al., 2015; Le Ven et al., 2016; Petersen et al., 2017), we noticed within the WT group a positive correlation between age and the volume of the OFT relative to the entire heart (% OFT volume; Figure 5C). Furthermore, retrograde OFT flow, which was present in two WT animals, correlated with atrial enlargement.

In addition to the 11 correlations observed in WT animals, we identified 37 new correlations in mutants (Figure 5D), suggesting a disruption of the morphology and/or function of the mutant heart. For example, we were able to address whether defects in OFT morphology affect cardiac performance and hemodynamics and vice versa. Indeed, we identified a negative correlation between OFT wall volume and heart rate, as well as between OFT luminal volume relative to the OFT (% OFT luminal volume) and OFT velocity time integral (VTI), suggesting that OFT morphology is linked with cardiac performance and blood velocity. We also observed a negative correlation between OFT volume and OFT wall volume with the regurgitation fraction in the AVC, suggesting that a more pronounced AVC retrograde flow is linked with defective (smaller and thinner) OFTs. Furthermore, there was a positive correlation between % OFT luminal volume and regurgitation fraction in the OFT, suggesting that larger OFT lumens are associated with higher retrograde flow in the OFT. When analyzing whether these morphological OFT defects were influenced by age or body size, we found that OFT total and wall volumes were negatively correlated with standard length and body volume, suggesting an aggravation of these phenotypes in older and bigger fish.

Regarding other cardiac compartments, the extent of retrograde flow across the AVC was associated with a smaller ventricle and bigger atrium, consistent with the consequences of valve regurgitation which leads to a higher amount of blood stalling in the atrium (Chaudhry et al., 2019). Interestingly, mutants - but not WT - with a higher regurgitation fraction in the OFT also displayed higher regurgitation fraction in the AVC, suggesting that these defects are closely associated.

In summary, these correlation analyses helped us identify links between functional and morphological phenotypes, even when statistical tests comparing WT and mutant groups failed to provide significance due to the high variability of the defects.

### Combined imaging analyses facilitate the selection of specific functional phenotypes for downstream morphological characterization

Having found that *alk5a* mutants frequently display retrograde blood flow and increased expansion of their OFT lumen, we analyzed additional animals using echocardiography and MRI to perform further analyses (Figure 6 A-E’). We chose an *alk5a*^*-/-*^ zebrafish that displays severe retrograde blood flow through the AV canal (Figure 6 A, D, E), OFT blood regurgitation (Figure 6B, D’, E’), as well as an unusually large expansion of the OFT lumen (Figure 6C). To analyze the morphological and ultrastructural defects in this animal, we performed light microscopy (Figure 6 F, G) and transmission electron microscopy (TEM; Figure 6 H-I’’) on tissue sections. Semi-thin sections for light microscopy showed a dilated lumen and thinner wall in the OFT (Figure 6F, G). TEM analysis revealed that the lumen of the WT OFT was barely detectable and lined with flattened or cuboidal endothelial cells (ECs; Figure 6H). We also observed that the WT OFT wall was composed of both spindle-shaped and cuboidal smooth muscle cells (SMCs; Figure 6H’, H’’) surrounded by a dense ECM. In contrast, the mutant OFT displayed an enlarged lumen lined by rounded ECs (Figure 6I). Notably, many cells lining the lumen in *alk5a*^*-/-*^ OFT appeared empty or detached, suggesting cell death. Despite a thinner OFT wall and a sparser and less dense ECM in the *alk5a*^*-/-*^ OFT, there were no obvious alterations in the SMCs (Figure 6I’, I’’).

**Figure 6.**
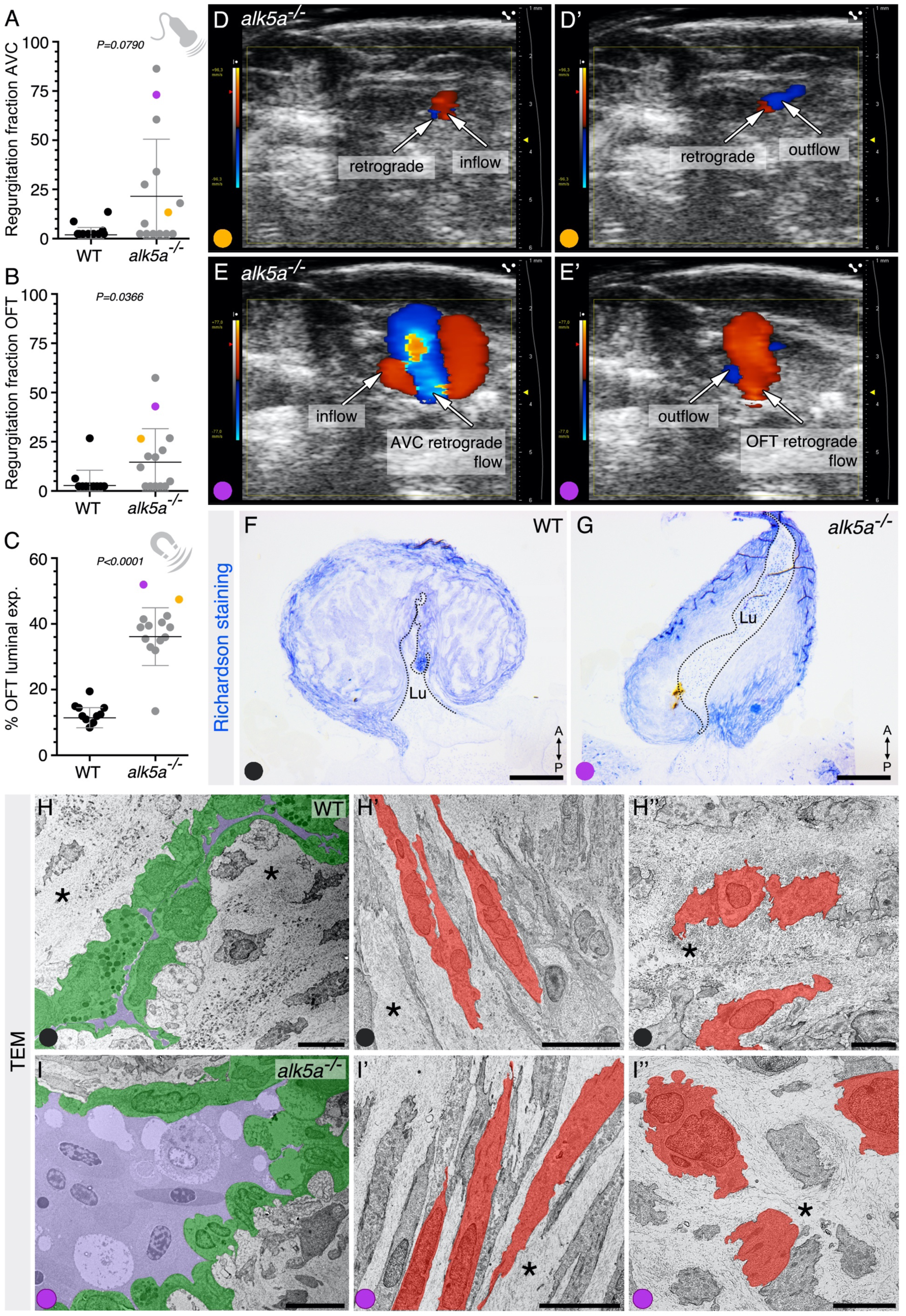
Combined imaging analyses facilitate the selection of specific functional phenotypes for high resolution morphological characterization. (A-C) Regurgitation fraction as obtained by echocardiography (A, B) and OFT luminal expansion as obtained by MRI (C) for two fish (orange and purple) plotted against the previously measured fish (black and gray). Plots show the values for each individual and the mean ± SD; *P*-values were determined by Mann-Whitney test. Purple dot refers to the fish selected for subsequent analysis with transmission electron microscopy (TEM). (D-E’) *alk5a*^*-/-*^ zebrafish showing mild (orange, D, D’) and severe (purple, E, E’) regurgitation fraction in the AVC and OFT regions. (F, G) Semi-thin sections of WT and *alk5a*^*-/-*^ OFTs stained with Richardson staining solution. Dashed line outlines the OFT lumen (Lu). (H-I’’) TEM images of WT and *alk5a*^*-/-*^ OFTs, showing the OFT lumen in purple, lined by ECs in green (H, I) and the OFT wall (H’, H’’, I’, I’’), including SMCs (red) and ECM (asterisk). Scale bars: 200 μm (F, G), 5 μm (H), 10 μm (H’-I’’).

Overall, these data indicate that non-invasive imaging techniques such as echocardiography and MRI represent useful platforms to identify individuals displaying functional phenotypes for further downstream high-resolution analyses.

## Discussion

In this study, we optimized and combined multiple cardiac imaging modalities including *in vivo* echocardiography and MRI along with *ex vivo* μ-CT to characterize the adult zebrafish heart (Figure 7). For each animal, we measured 32 different cardiac parameters in WT and *alk5a* mutant samples. Bulk analyses of mutants vs. WT, as well as intra-individual correlations between these parameters led to the identification of a primary OFT phenotype.

**Figure 7.**
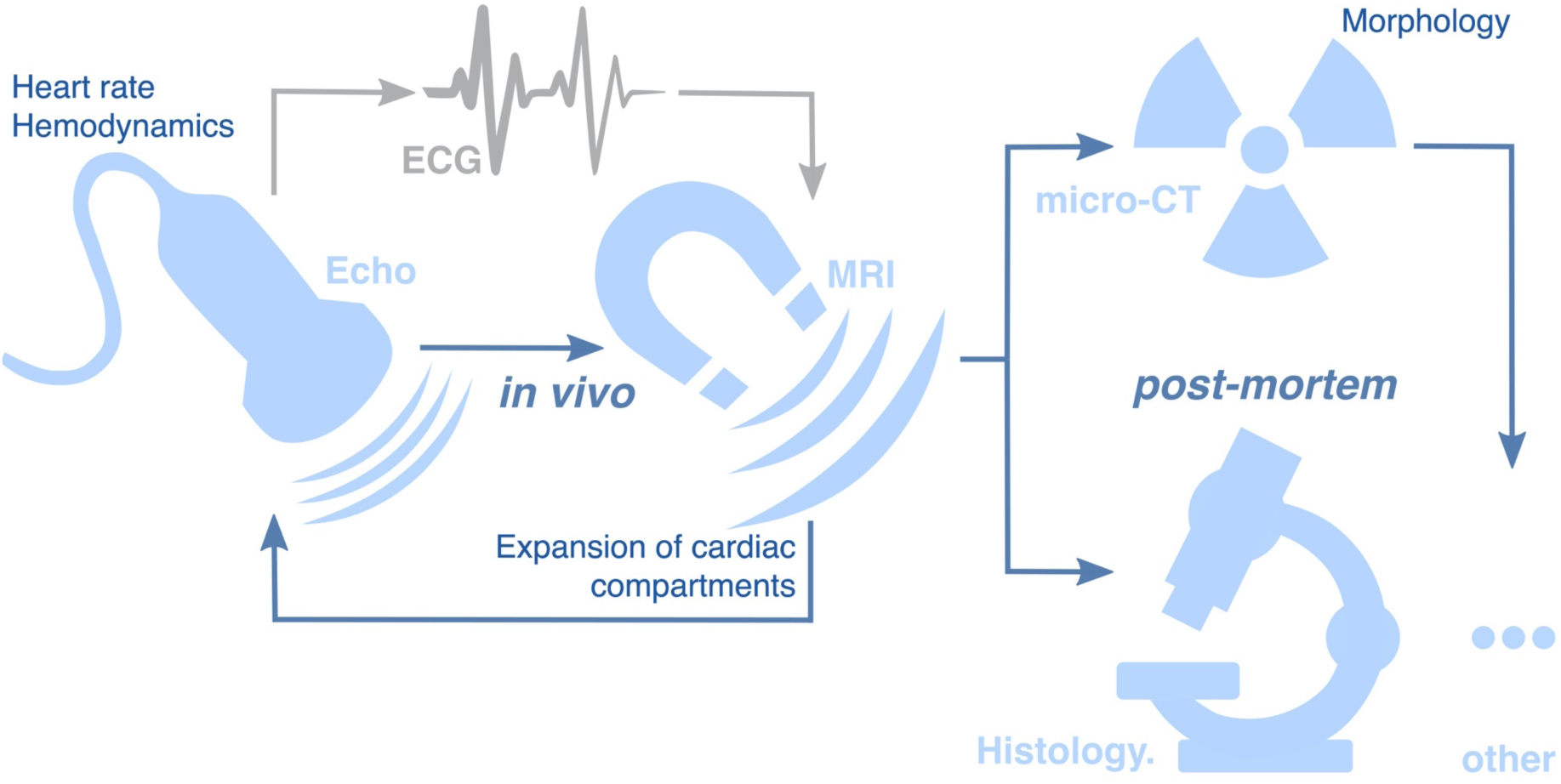
Schematics of a multimodal preclinical imaging platform for adult cardiac phenotyping. Echocardiography allows for *in vivo* characterization of heart rate and hemodynamics; MRI allows for analysis of the expansion of the cardiac compartments in virtual reconstructions of beating hearts. These imaging analyses can be used for longitudinal studies and can be complemented with cardiac ECGs to determine the electrophysiological effects. Morphological analyses can be performed with μ-CT imaging to determine alterations in cardiac volumes and anatomy. In parallel, low-throughput analyses such as histology and molecular profiling can be done to better characterize the cardiac phenotypes identified with the multimodal imaging approach

### Optimization of preclinical cardiac imaging analysis

While during development it is possible to use microscopy for single-cell resolution imaging of the zebrafish internal organs, as animals grow, the tissues become opaque forcing researchers to rely on tissue sections. Previous publications have reported the optimization of preclinical imaging techniques such as echocardiography, static MRI, and μ-CT for adult zebrafish. However, the zebrafish field lacks reference values for each technique, methods to reconstruct beating hearts imaged with MRI, and protocols to analyze data from different techniques.

Here, taking advantage of the echocardiography Color Doppler Mode, we were able to gather data on the hemodynamics within the heart and aorta in adult zebrafish. We observed blood flow perturbation in some *alk5a* mutant fish including blood regurgitation in the AV and OFT canals, and increased aortic flow.

With our adapted MRI protocol, we were able to observe the contracting heart with considerably improved image contrast and resolution, compared with echocardiography. This technique allowed us to define the cardiac performance in adult fish and to observe that 11/12 *alk5a*^*-/-*^ animals exhibited a severely expanded OFT.

We complemented this functional analysis with ex-vivo μ-CT imaging to examine the morphology of the adult zebrafish heart in fine detail. With a few exceptions (Descamps et al., 2014; Vagberg et al., 2015; Babaei et al., 2016; Weinhardt et al., 2018; Ding et al., 2019), this technique has been mainly used for ex-vivo samples focusing on the skeleton (Charles et al., 2017; Hur et al., 2017; Silvent et al., 2017; C. J. Watson et al., 2020). 3D rendering of the cardiac compartments and aorta allowed the identification of alterations specifically in the OFT, atrial volume, and diameter of the aorta in some *alk5a* mutant fish. Since this imaging modality does not require tissue dissection, the preservation of tissue integrity and organ positioning brings additional information, allowing the investigation of phenotypes affecting the great vessels and other connecting structures. Further development of μ-CT for *in vivo* imaging will have to take into consideration the exposure to radiation and the need to immobilize the sample during data acquisition. This approach would be especially relevant to study the progression of conditions affecting cardiac performance, including during cardiac regeneration.

Moreover, as clinicians strongly rely on reference values to define a healthy heart considering the gender, age and ethnicity of the patients (Lancellotti, 2014; Caballero et al., 2015; Kawel-Boehm et al., 2015; Le Ven et al., 2016; Petersen et al., 2017), we define some reference values for each technique (Tables 3-5) which will help future studies.

### Multimodality approach

When designing studies with adult specimens, the reduced sample size and the high individual variability need to be considered. Moreover, the same disease-causing mutation can lead to a phenotype in only some of the mutant individuals (incomplete penetrance), or lead to different phenotypic severity (variable expressivity) (Cooper et al., 2013). In our analyses, we observed high variability among individuals, particularly related to hemodynamic measurements. Overall, we observed that few of the parameters measured with each technique were significantly different (defined as a *p*-value <0.05) between WT and mutants. Nevertheless, it was evident that some *alk5a* mutant fish presented cardiac defects. Therefore, in order to investigate this variability between individuals, we subjected the same animal to multiple imaging techniques to increase the data collected per individual and decrease the number of specimens required. We used these techniques to select specimens with specific features for high resolution approaches including TEM. This approach is similar to that proposed by clinicians who have advocated the use of multimodal imaging approaches for a more accurate diagnosis of complex and variable cardiac diseases (Kirisli et al., 2012; Cardim et al., 2015; Galderisi et al., 2015; Plana et al., 2018).

In zebrafish thus far, only a few studies have integrated different modalities for phenotypic characterization, for example combining μ-CT and electron microscopy (Silvent et al., 2017; De Spiegelaere et al., 2020). Here, we analyzed parameters that encompass hemodynamics, cardiac performance, and morphology in the same fish. It will be interesting to complement this multimodal platform with electrocardiography to obtain data on the electrophysiology of the heart (Dhillon et al., 2013; Liu et al., 2016; Lenning et al., 2017; Ahlberg et al., 2018; Le et al., 2019; Zhao et al., 2019) (Figure 7). Moreover, by correlation analysis of each parameter, in addition to bulk analysis of WT vs mutant samples, and by selecting individuals with specific functional phenotypes, we achieved a better understanding of the cardiac phenotypes.

### Clinical perspective of the *alk5a* cardiac phenotype

Overall, our data indicate that while the loss of *alk5a* alone does not cause embryonic defects (Boezio et al., 2020), it can cause a cardiac defect symptomatic only at later stages. Our data also identify the OFT (orthologous to the mammalian aortic root) as the main compartment affected in *alk5a* mutants. Interestingly, many of the phenotypes we observed are similar to defects associated with aortopathies in clinical settings, pathologies often derived from mutations in TGF-β pathway genes (Loeys et al., 2006; Matyas et al., 2006; Camerota et al., 2019). At the morphological level, we observed that *alk5a* mutants present a thinning of the OFT outer wall, similar to patients affected by aortic aneurysms prone to dissection (Van Puyvelde et al., 2016; Jana et al., 2019). It is also interesting to note that the volume of the OFT wall decreased according to the age and size of the fish, similar to the decrease in aortic wall thickness in older patients (Fhayli et al., 2019; Clift et al., 2020). At the functional level, we observed that increased OFT dilation was linked with retrograde blood flow, as often reported in patients presenting aortic dilation and regurgitation (Roman et al., 1987; Nataf et al., 2006; Tadros et al., 2009; Evangelista et al., 2010). In addition, the increased aortic flow phenotype in *alk5a* mutant fish resembles what has been reported in human patients, in whom aortic dilation can cause disturbed and tortuous flow not only in the aneurysm site but also in apparently unaffected aortic portions (Numata et al., 2016).

In summary, we propose a combinatorial imaging and data analysis platform to detect phenotypes which are too variable or subtle to identify with conventional histological analyses. This study further demonstrates the importance of the zebrafish model to clarify adult phenotypes potentially translatable to the clinic.

## Materials and Methods

### Zebrafish husbandry and lines

All zebrafish husbandry was performed under standard conditions, and all experiments were conducted in accordance with institutional (MPG) and national ethical and animal welfare guidelines. All the fish used in the study were in the *Tg(kdrl:eGFP)*^*s843*^ (Jin et al., 2005) background.

The *tgfbr1a* ^*bns329*^ fish (Boezio et al., 2020) carry a 2 bp insertion and a 6 bp deletion in exon 1. For all the experiments, maternal zygotic *tgfbr1a*^*-/-*^ fish were analyzed and compared to WT fish. The mutants were genotyped as previously described, using high-resolution melt analysis (HRMA) (Eco-Illumina) and/or sequencing with the following primers: *alk5a* HRM fw: 5’-CTTCTGGACAGACCGTGACA-3’, *alk5a* HRM rv: 5’-GAAGGAGCGCACTGGAAAG -3’.

### Confocal microscopy for live imaging of zebrafish larvae

For live confocal imaging, embryos and larvae were embedded in 1% low-melting agarose/egg water with 0.2% tricaine (Sigma) to image stopped hearts. Larvae were imaged with LSM800 Axio Examiner confocal microscope with a W Plan-Apochromat 40×/1.0 or W Plan-Apochromat 20×/1.0 dipping lenses. In order to image the heart, the larvae were placed in a supine position. All images were acquired using the ZEN Blue (Zeiss) software.

### Histology and Immunostaining

Adult hearts were fixed in 4% buffered paraformaldehyde for 1 hour at room temperature, washed in 1x PBS and embedded as previously described (Mateus et al., 2015). Briefly, the tissue was placed overnight at 4°C in a solution of 30% (w/v) sucrose prepared in 1x PBS, pre-embedded in 7.5% (w/v) porcine gelatin (Sigma)/15% (w/v) sucrose in 1x PBS at 37°C for 1 hour and embedded with a new solution of gelatin. Tissue blocks were frozen in isopentane (Sigma) cooled in liquid nitrogen. Cryosections were cut at 10 μm using a Leica CM3050S cryostat (Leica) and kept at −20°C until further use. Prior to further analysis, slides were thawed for 10 minutes at room temperature and gelatin was removed in 1x PBS at 37°C. For hematoxylin and eosin staining, cryosections were stained with acidic hemalum (Waldeck) for 10 minutes, washed in running tap water for 2 minutes and rinsed in deionized water. The sections were then counterstained with eosin (Waldeck) for 6 min, dehydrated in 100% ethanol, cleared in xylene and mounted in entellan (Merck). Sections were imaged using a Nikon SMZ25 microscope.

Immunostaining started with a wash in 0.1M glycine (Sigma) followed by permeabilization for 7 minutes at −20°C in pre-cooled acetone. Sections were incubated in a blocking solution of PBDX (1% (w/v) Bovine Serum Albumin, 1% (v/v) DMSO, 1% (v/v) Triton-X100 in PBS) with 15% (v/v) goat serum for a minimum of two hours at room temperature.

Incubation with the following primary antibody was performed overnight at 4°C: anti-GFP chicken (Aves Technology; 1:200), anti-Elastin2 rabbit (Miao et al., 2007) (1:100). Slides were washed several times with PBDX and incubated with the corresponding secondary antibodies (1:500) overnight at 4°C: anti-chicken AlexaFluor 488 (ThermoFisher), anti-rabbit AlexaFluor647 (ThermoFisher). For all incubations, slides were covered with a piece of Parafilm-M to ensure homogenous distribution of the solution. Slides were washed a minimum of 3 times for 15 minutes each in a solution of 0.3% (v/v) Triton-X100 in PBS (PBST) and counterstained with 0.0002% (w/v) DAPI (Merck) in PBST for 10 minutes. Slides were then washed a minimum of 3 times for 15 minutes each in PBST and mounted with DAKO Fluorescence mounting medium (Agilent). Elastin2 antibody was purified from the previously described serum stock (Miao et al., 2007). Sections were imaged using a Zeiss LSM700 (Zeiss) microscope, and ZEN software (Black edition, Zeiss).

### Transmission Electron Microscopy (TEM)

Adult hearts were dissected and fixed in 4% PFA with 2.5% glutaraldehyde in 0.05 M HEPES buffer (pH 7.2) for 2 hours at room temperature, and subsequently stored at 4°C. Samples were rinsed three times in 0.05 M HEPES buffer (pH 7.2) and post-fixed in 1% (w/v) OsO_4_ for 1 hour. After washing three times with distilled water, blocs were stained with 2% uranyl acetate for 1 hour. Samples were dehydrated through a graded series of ethanol washes, transferred to propylene oxide, and embedded in Epon according to standard procedures (Laue, 2010). Tissue semi-thin sections (900 nm thick) were obtained in a Ultracut E microtome (Reichert-Jung, Leica) and stained with Richardson staining solution (Richardson et al., 1960). Ultra-thin 70 nm sections were then collected on copper grids. After post staining with uranyl acetate and lead citrate, ultrathin sections were examined with a JEM-1400 Plus transmission electron microscope (Jeol, Japan), operated at an accelerating voltage of 120 kV. Digital images were recorded with an EM-14800Ruby Digital CCD camera unit.

### Body volume measurements

Total body volume was determined by submerging each animal in a 2 mL syringe filled with water and determining water displacement. 1 mL water displacement corresponds to 1 cm^3^ of body volume.

### Doppler Echocardiography

Zebrafish were anesthetized using 0.016% buffered Tricaine diluted in system water. Animals were placed in supine position in a bed made of modeling clay, adjustable to the size of the fish, and submerged in anesthesia solution to ensure the propagation of the ultrasound signal. Vevo2100® Imaging System (VisualSonics) and VisualSonics Ultrasound Imaging Software (Version 1.6.0) were used for echocardiography. Since the required ultrasound frequency depends on the size of the sample (Benslimane et al., 2019), we used a high-frequency MicroScan transducer (MS700 v3.0) at a frequency of 40 MHz, as opposed to the 15 MHz used clinically for humans. The transducer was oriented along the longitudinal axis of the fish and the images were acquired with the anterior side of the fish to the left and posterior side to the right (Figure 2 - figure supplement 1 A). The imaging was performed in Color Doppler mode. Videos were recorded for 10 seconds and across different 2D planes, spanning atrium, ventricle and OFT. Image acquisition was completed within 5 minutes of sedation to avoid cardiac function aberrations. After imaging, the fish were returned to system water and observed until recovery. Vevo Lab™ software package v.1.7.0 (VisualSonics) was used for image analysis.

The heart rate was calculated as beats per minute in the 10 second time frame recorded. The inflow and outflow area were measured in the 2D planes exhibiting the highest blood velocity across the heart. The regurgitation fraction across the AVC or OFT were calculated as the ratios between the area of the inflow and outflow present simultaneously in the same plane spanning the AV or OFT canal, respectively. The area of the aortic flow refers to the area of flow rostral to OFT, if detected. Values as VTI, mean and peak gradient, mean and peak velocity and ejection time were calculated automatically by Vevo Lab™ software package v.1.7.0 (VisualSonics) and refer to the curves highlighted in Figure 2- figure supplement 1B, B’.

### Magnetic Resonance Imaging (MRI)

Cardiac MRI measurements were performed on a 7.0 T Bruker PharmaScan (Bruker Biospin, Ettlingen, Germany) small animal MRI system. The instrument was equipped with a 780 mT/m gradient system, a cryogenically cooled 4 channel phased array element ^1^H receiver-coil (CryoProbe™, Bruker Biospin) and a 72 mm room temperature volume resonator for transmission (Bruker Biospin). Zebrafish were anesthetized in 0.016% Tricaine solution (Sigma) and placed in a supine position in a bed made of modelling clay. We decided to analyze our specimens in a container without water flow and reduce the imaging time to less than 20 minutes to ensure animal survival.

Localizer images were acquired using a spin-echo sequence and correction of slice orientation was performed when necessary. In addition, RARE (Rapid Acquisition with Relaxation Enhancement) sequences (TR=2500 ms, TE=36.7 ms, slice thickness=0.3 mm) in a sagittal orientation were used to determine the correct coronal plane for imaging the ventricle or OFT (Figure 3 – figure supplement 1 A-B’). The imaging was performed with the following parameters: IntraGate 2D FLASH: TE/TR: 4.2/8.2 ms, flip angle 10°, slice thickness: 0.3 mm, in-plane resolution: 59 × 59 or 47 × 47 µm^2^. Navigator based self-gating (Echteld et al., 2006; Nauerth et al., 2006) was used to determine the cardiac rate and performed cine retrospective reconstruction. With the help of the profile images and the navigator signal, we were also able to detect and exclude sudden movements of the anesthetized fish as well as shifts in resting position (Figure 3 – figure supplement 1 C-C’’).

The evaluation of the functional heart data of adult zebrafish, following MRI, was performed with the Qmass® MR 8.1 (Medis Medical Imaging Systems) software. The percentage of OFT and ventricular expansion was calculated by the software after manually defining the lumen (or wall) outline during maximum and minimum chamber dilation. The OFT and ventricular measurements were performed in different recordings with different orientations selected to show the cross-section of the desired chamber. All the values represent the average of two independent analyses of the same data performed by two different scientists.

### Micro-CT

After the MRI recordings, the fish were overdosed with anesthetic, fixed in ice-cold 4% PFA and kept in PFA over-night at 4°C. As previously described (Descamps et al., 2014), a solution of 2.5% phosphomolybdic acid (PMA) prepared in demineralized water was used to stain each fish individually for a period of 6 days. Samples were then gradually transferred to a 70% ethanol solution where they were preserved at room temperature until imaging.

All samples were scanned by the µ-CT scanner model Skyscan 1276 (Bruker). X-ray parameters for the imaging were: Source voltage = 55kV; source current = 72 µA; image pixel size = 2.965 µm; filter = Al 0.25 mm; rotation step = 0.25 deg to 180 deg. For image reconstruction, we used NRecon (version 1.7.1.0) and InstaRecon (version 2.0.3.7) softwares. Resulting reconstructed images were 3856 × 2248 pixels, with a pixel size of 2.965 µm. All the morphological measurements were performed with Imaris (Bitplane) software. The atrium, ventricle, OFT and aorta were manually segmented across the entire 3D volume. The lumen of the OFT was also separately segmented. The volumes of each compartment were calculated with the “Surface tool” in Imaris and, when required, normalized to the heart volume (Figure 4-figure supplement 1 C-E) or total OFT volume (Figure 4H). The OFT wall volume was calculated by subtracting the OFT luminal volume from the total OFT volume. The diameter of the aorta was consistently measured in the most proximal third of the aorta.

### Data analyses and quantification

All statistical analyses were performed in GraphPad Prism (Version 6.07) and illustrations were done in Inkscape (XQuartz X11).

For comparison of WT vs mutant samples, Gaussian distribution was tested for every sample group using the D’Agostino-Pearson omnibus normality test. When both samples passed the normality test, *P*-values were determined by unpaired *t-*test. When, at least one of the samples did not pass the normality test, *P*-values were determined using the Mann-Whitney test for comparison of 2 samples.

To address the linear relationship between all possible pairs of parameters, correlation matrices were generated through the calculation of Pearson’s correlation coefficients. A binary numerical system was used to compute the sex of individuals (0, male; 1, female). Calculations and corresponding correlograms were generated using RStudio v1.1.456 (RStudio Team, 2015), and the *Hmisc* (v 4.4-0; Harrel Jr, 2020) and *corrplot* (v0.84; Wei, 2017) packages. The significance level was set to α = 0.05 for all bivariate correlations.

## Acknowledgments

We would like to thank Dominique Adriaens for insights regarding the µ-CT contrast agent protocol; Simon Perathoner, Srinath Ramkumar and Ursula Hofmann for assistance; Dimitris Beis for the Elastin 2 antibody; Jesper Hastrup for valuable discussion; all the fish facility staff for technical support; and Matteo Perino, Pia Lundegaard, Michelle Collins, Samuel Capon, Felix Gunawan, Rubén Marín-Juez, Margareta Albu and Mridula Balakrishnan for critical comments on the manuscript. Research in the Stainier Lab is supported in part by the Max Planck Society, the DFG (Sonderforschungsbereich - SFB 834), the Leducq Foundation and the European Research Council (AdG project: ZMOD 694455).

## Author contributions

Conceptualization, A.B.B., G.L.M.B. and D.Y.R.S.; Methodology, A.B.B, G.L.M.B, A.W., R.R., J.P.; A N. and C.M.; Validation, A.B.B., G.L.M.B., J.C.S.; Formal Analysis, A.B.B, G.L.M.B., J.C.S.; Investigation, A.B.B., G.L.M.B., A.W., J.P.; Resources, C.S.M.H; Writing – Original Draft, A.B.B., G.L.M.B. and D.Y.R.S.; Writing – Reviewing & Editing, all; Visualization, A.B.B, G.L.M.B.; Supervision, A.B.B. and D.Y.R.S; Project Administration, A.B.B. and D.Y.R.S.; Funding Acquisition, D.Y.R.S.

---

**Figure 1 - figure supplement 1 – *alk5a* mutant larvae exhibit no obvious morphological defects**.

(A, B) Brightfield images of 96 hpf WT (A) and *alk5a*^*-/-*^ (B) larvae show no obvious morphological differences. (C-D’) Confocal images of 96 hpf WT (C) and *alk5a*^*-/-*^ (D) *Tg(kdrl:eGFP)* hearts. Boxed area is shown in C’ and D’. A- atrium, V- ventricle, OFT- outflow tract. Scale bars: 400 μm (A, B), 50 μm (C, D), 20 μm (C’, D’).

**Figure 2 - figure supplement 1 – Most parameters obtained with echocardiography do not show differences between WT and mutant zebrafish**.

A) Schematic of the echocardiography settings, depicting the position of the transducer relative to the heart (red arrow, AVC blood inflow; OFT blood outflow). (B-C’) Quantification of hemodynamic parameters, mostly focused on the OFT flow (blue, B; negative curves, C and C’). (D-K) Plotted values for WT and *alk5a*^*-/-*^ adult zebrafish obtained with Doppler echocardiography analyses. The color of each dot refers to the same fish across all graphs. The number symbol refers to the individual fish mentioned in the text. Plots show the values for each individual and the mean ± SD; *P*-values were determined by unpaired *t-* test (D-I, K) or Mann-Whitney test (J). AU – arbitrary units.

**Figure 3 - figure supplement 1 – MRI analyses of beating hearts do not show differences in ventricular expansion**.

(A-B’) Examples of RARE (Rapid Acquisition with Relaxation Enhancement) images of a female (A, A’) and male (B, B’) WT adult zebrafish; sagittal views. Dashed lines define the orientation chosen to image OFT (A, B) or ventricular (A’, B’) expansion. (C-C’’) Graphs showing raw signal including heartbeat and sudden movements of the fish (arrows, C) and filtered signal excluding fish movements (C’) to accurately determine heart rate (C’’). Boxed area is shown in C’’. (D, E) Percentage of ventricular outer wall (D) and luminal (E) expansion in WT and *alk5a*^*-/-*^ adult zebrafish. Plots show the values for each individual and the mean ± SD; *P*-values were determined by unpaired *t-*test (D) or Mann-Whitney test (E). (F-G’’) Single frames of MRI cines of WT (F) and *alk5a*^*-/-*^ (G) zebrafish in coronal view. Boxed area is shown in F’, G’ (systole) and F’’, G’’ (diastole). Magenta dashed line, ventricular lumen; yellow dashed line, ventricular outer wall. The color of each dot refers to the same fish across all graphs and images. The number symbol refers to the individual fish mentioned in the text. A- atrium, V- ventricle, OFT- outflow tract. Scale bars: 1.5 mm (C, D), 500 μm (C’, C’’, D’, D’’).

**Figure 4 - figure supplement 1 – All cardiac compartments display variable volumes in *alk5a* mutants**.

(A-A’’) Orthogonal views of a WT adult zebrafish imaged with μ-CT showing coronal (A), axial (A’) and sagittal (A’’) views. (B-G) Quantification of morphological parameters for each cardiac compartment in WT and *alk5a*^*-/-*^ adult zebrafish. Plots show the values for each individual and the mean ± SD; *P*-values were determined by unpaired *t-*test (B, E-G) or Mann-Whitney test (C, D). The color of each dot refers to the same fish across all graphs.

The number symbol refers to the individual fish mentioned in the text. Scale bars: 1 mm (A-A’’).

**Video 1 –** Doppler Echocardiography of a WT fish exhibiting separate blood inflow (red) and outflow (blue), without signs of regurgitation. Related to Figure 2E, E’.

**Video 2 –** Doppler Echocardiography of an *alk5a*^-/-^ fish exhibiting blood flow regurgitation in the AVC and aortic flow directly rostral to the heart. Related to Figure 2F, F’.

**Video 3 –** Doppler Echocardiography of an *alk5a*^-/-^ fish exhibiting severe blood flow regurgitation in the AVC and OFT. Related to Figure 2G, G’.

**Video 4 –** MRI imaging of a WT fish in coronal view showing ventricular expansion. Related to Figure 3- figure supplement 1 F-F’’.

**Video 5 –** MRI imaging of an *alk5a*^*-/-*^ fish in coronal view showing ventricular expansion. Related to Figure 3- figure supplement 1 G-G’’.

**Video 6 –** MRI imaging of a WT fish in coronal view showing OFT expansion. Related to Figure 3C-C’’.

**Video 7 –** MRI imaging of an *alk5a*^*-/-*^ fish in coronal view showing OFT expansion. Related to Figure 3D-D’’.

**Video 8 -** µ-CT scans of the anterior region of a WT fish in sagittal view. Related to Figure 4 A.

**Video 9 -** µ-CT scans of the anterior region of an *alk5a*^*-/-*^ fish in sagittal view. Related to Figure 4 B.

**Video 10 –** Volumetric surface rendering of the cardiac compartments (blue, atrium; red, ventricle; grey, OFT; purple, OFT lumen) of a WT fish imaged with µ-CT. Related to Figure 4C, C’’.

**Video 11 –** Volumetric surface rendering of the cardiac compartments (blue, atrium; red, ventricle; grey, OFT; purple, OFT lumen) of an *alk5a*^*-/-*^ fish imaged with µ-CT. Related to Figure 4D, D’’.

**Video 12 –** Volumetric surface rendering of the OFT (grey), OFT volume (purple) and aorta (pink) of a WT fish imaged with µ-CT. Related to Figure 4 C’.

**Video 13 –** Volumetric surface rendering of the OFT (grey), OFT volume (purple) and aorta (pink) of an *alk5a*^*-/-*^ fish imaged with µ-CT. Related to Figure 4 D’.

**Video 14–** Volumetric surface rendering of the OFT (grey), OFT volume (purple) and aorta (pink) of a WT fish imaged with µ-CT, as seen in a transversal section starting from the OFT and proceeding rostrally. Related to Figure 4 C’’’’.

**Video 15 –** Volumetric surface rendering of the OFT (grey), OFT volume (purple) and aorta (pink) of an *alk5a*^*-/-*^ fish imaged with µ-CT, as seen in a transversal section starting from the OFT and proceeding rostrally. Related to Figure 4 D’’’’.

