## Supplemental Figures for "Integration of multiple imaging platforms to uncover cardiac defects in adult zebrafish"

**Figure 1 – figure supplement 1**

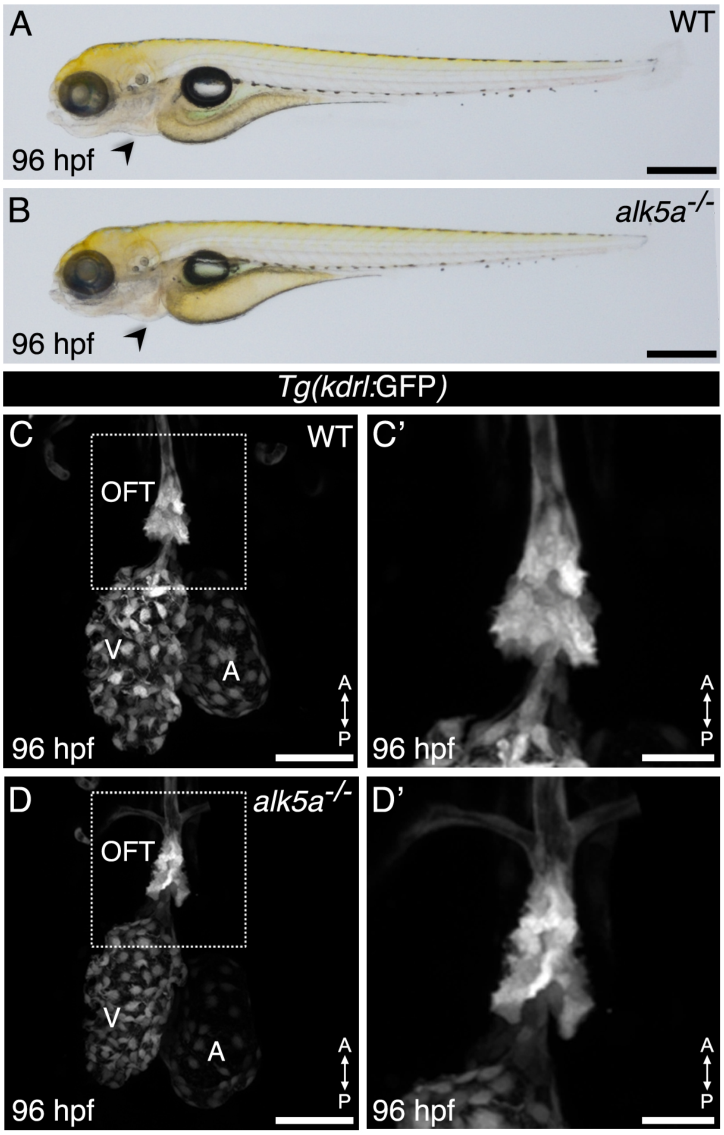

### Figure 2 – figure supplement 1

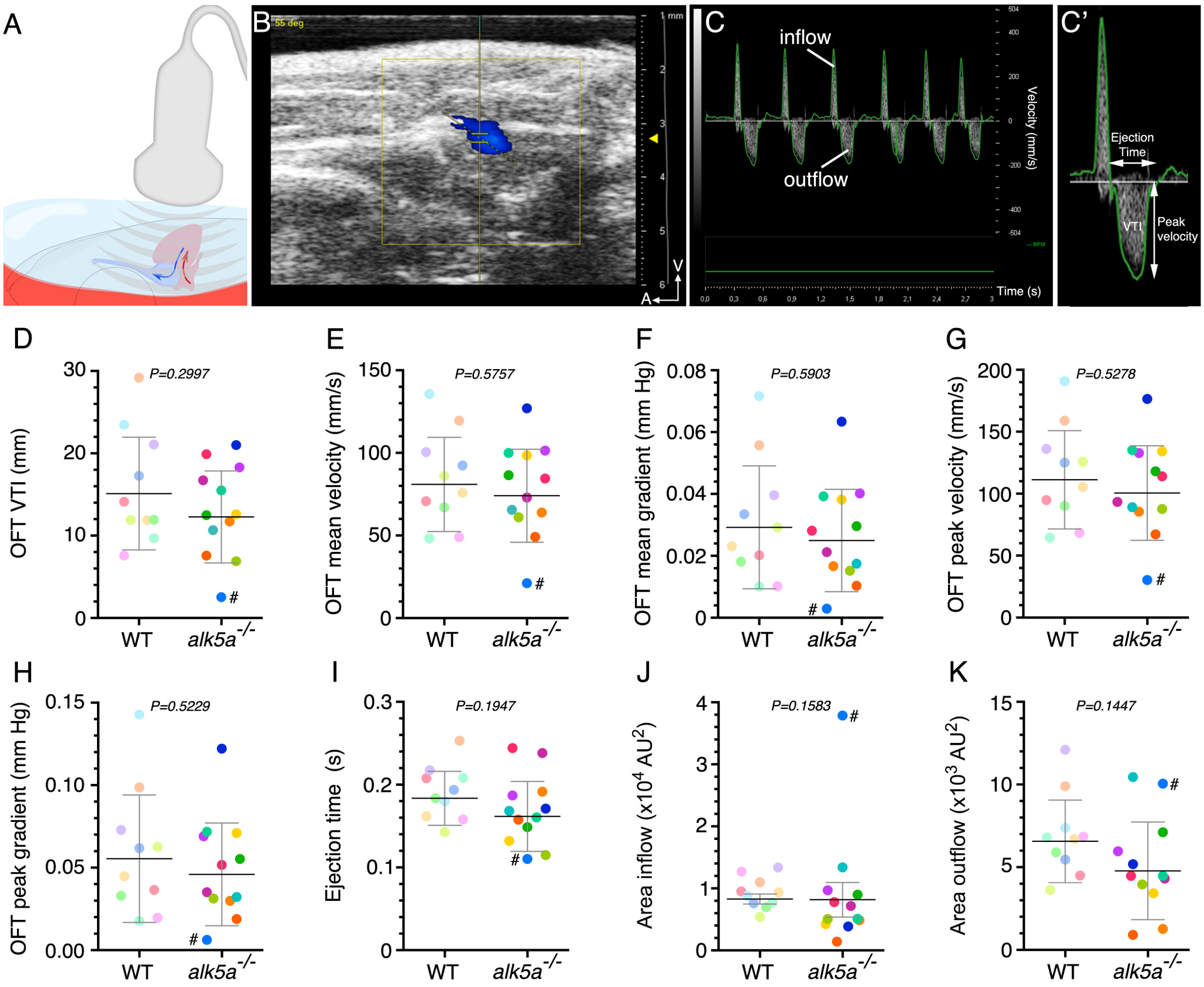

#### Figure 3 – figure supplement 1

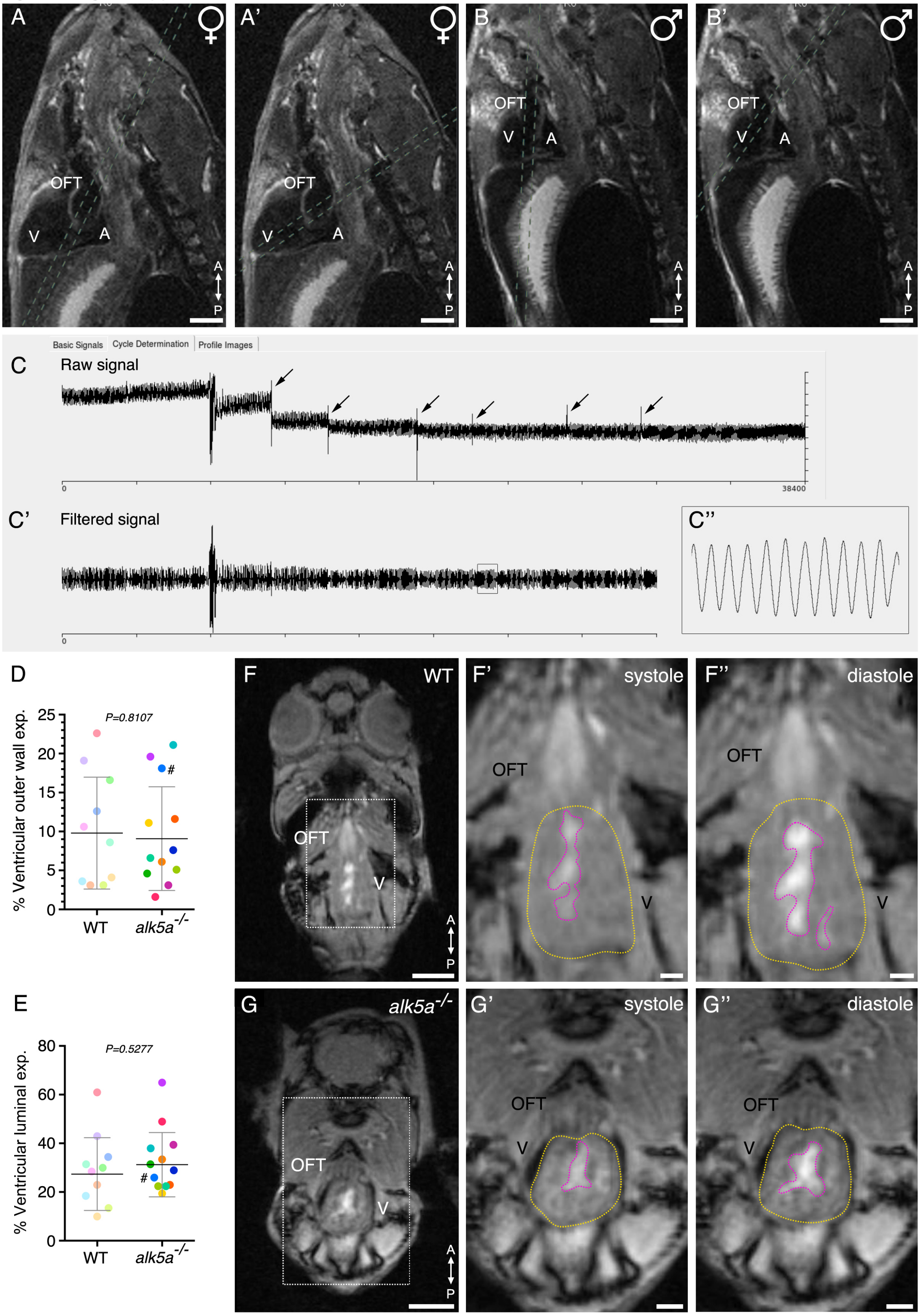

### Figure 4 – figure supplement 1

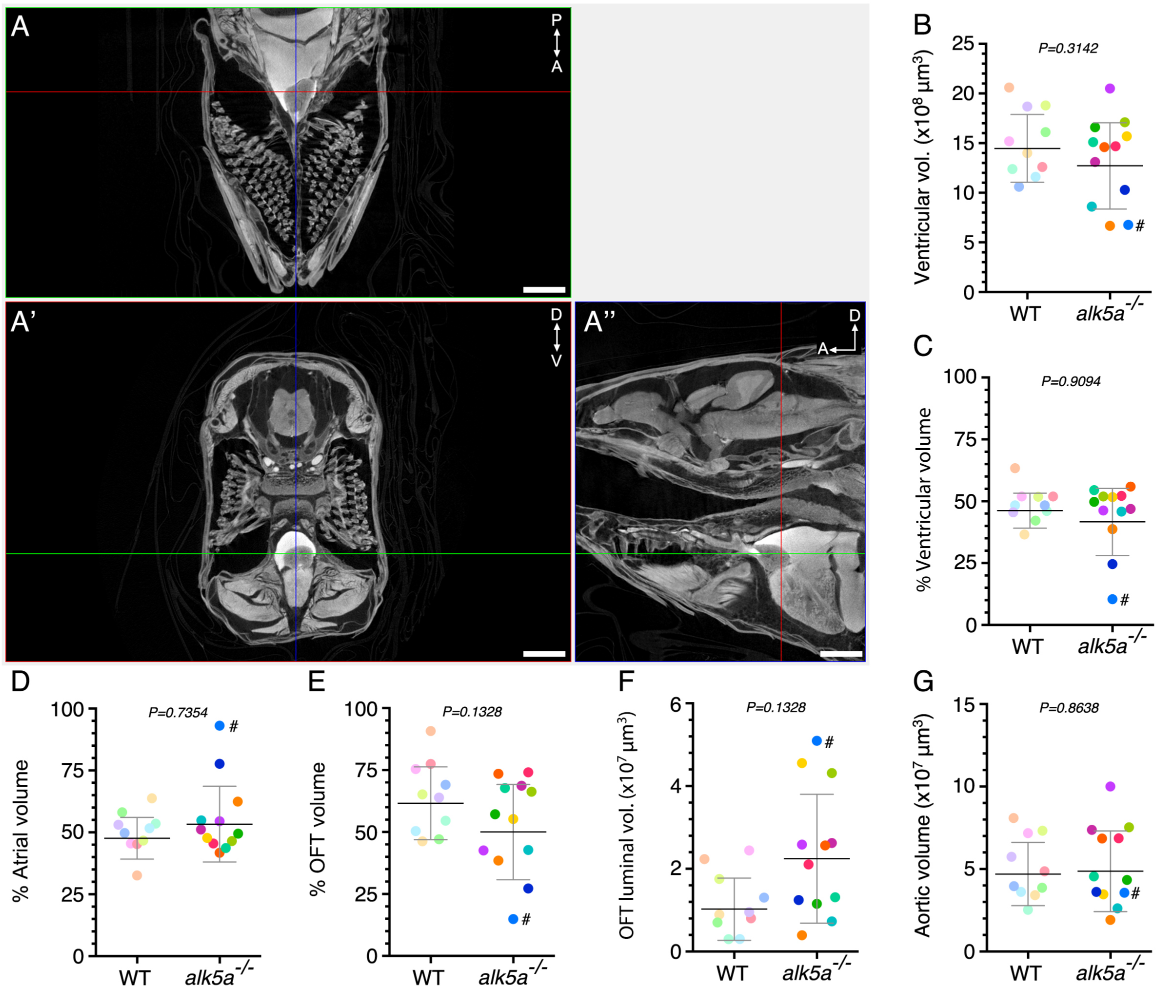
